# Improved subspecies identification in clinical *Mycobacterium abscessus* complex isolates using whole genome sequence

**DOI:** 10.1101/2020.01.02.893412

**Authors:** Dachuan Lin, Ruifeng Sun, Yaoju Tan, Jing Wang, Xinchun Chen

## Abstract

*Mycobacterium abscessus* complex, which is frequently reported causing a variety of skin and soft tissues diseases in humans, is composed of three subspecies, namely *M. abscessus subsp. abscessus, M. abscessus subsp. massiliense* and *M. abscessus subsp. bolletii*. Currently, the differentiation of these three subspecies in clinical isolates still largely depend on single gene identification methods like the genes namely *hsp65, 16s* with a limited accuracy. This study confirmed the limitations of the single gene based method in the subspecies identification. We performed a comprehensive analysis of MABC genomes in the NCBI database and tried to build an accurate and user-friendly identify method. Here, we describe an improved assay for *Mycobacterium abscessus* complex fast identification using WGS data, based on the identities of *rpoB, erm*(41) and *rpls*. Comprehensive analysis has been performed to compare our software results with the traditional method. The result showed that the method built-in this study could 100% identification the subspecies for the *Mycobacterium abscessus* complex in the public genome database (893 genomes from NCBI database and 6 clinical isolates from this study). Because this software can be easily integrated into a routine workflow to quickly and precisely provide subspecies-level identification and discrimination MABC different subspecies in clinical isolates by WGS. This assay will facilitate accurate molecular identification of species from the MABC complex in a variety of clinical specimens and diagnostic contexts.

## Introduction

*Mycobacterium abscessus* complex (MABC) which caused a range of diseases from skin infections to pulmonary [1, 2] is a rapidly growing mycobacterium which becomes an emerging pathogen. MABC is notorious not only because it can accelerate inflammatory damage which leading to increased morbidity and mortality, but also because it causes cystic fibrosis (CF) that had become the most lethal and frequent infection. The notorious MABC complex caused diseases were called nightmares in clinical field, not only because *M. abscessus* is the second most common nontuberculous mycobacterial species associated with lung disease, but also the intrinsic and acquired resistance of *Mycobacterium abscessus* to commonly used antibiotics limits the chemotherapeutic options for infections caused by these mycobacteria. So the infections caused by MABC especially the multidrug resistance strains were very difficult to treat [3-7], and sometimes impossible to treat, even in the developed countries [8, 9].

Despite the high genome similarity, the members of the MABC usually have distinct phenotypes in culture, antibiotic resistance pattern, particularly the critical first line antibiotic treatment-clarithromycin[10, 11], and especially cause differing treatment outcomes for patients infected with *M. abscessus subsp. abscessus* versus *M. abscessus subsp. massiliense*[12]. Different treatment requirements and outcomes are thought to vary among the different subspecies[13], thus it is clinically significant to differentiate MABC. So the main purpose of this study was to construct an accuracy and user-friendly method for MABC subspecies identification.

The MABC represent a diverse and clinically important family of bacteria. Although controversy still exists about the taxonomy and nomenclature of *M. abscessus* subspecies, most of the researchers believed that MABC comprised three subspecies: *M. abscessus subsp. abscessus, M. abscessus subsp. massiliense* and *M. abscessus subsp. bolletii*[14, 15].With the technical improvement, the debates (re-classification, elevated to species level) of classification of *Mycobacterium abscessus* had companied the discovery since 1953[16], even during the past 30 years the MABC taxonomic classification still have a serial changes from single species to different species then to three subspecies[17]. And the debate still last nowadays[18], so one of the purposes of this study was to further clarify the debate by the genome comparison.

Prior work had demonstrated the commonly used 16S rRNA was not good enough to identify MABC[19], and only using *rpoB* also could lead misidentifying for Mycobacterium[23]. Thus, in this study we evaluated the current taxonomic methods based on 16S rRNA, *rpoB, erm*(41) and *erm*(42) and in order to find a good MABC taxonomic classication method by the combination of those marker genes together. Another thing is as we all known assessing species boundaries among different species or subspecies is critical for taxonomy, thus special cut-off value is needed for different species, even for the traditional identification genes. So this study comprehensive compared the genome in database and selected the cut-off for the MABC taxonomic calssication. Additionally, to our knowledge, no specific genes were identified to discriminate *M. abscessus subsp. bolletii* (The traditional *rpoB* gene is not good enough for subspecies identification). To our knowledge, WGS-based protocols have so far not been developed for subspecies level like MABC identification. Therefore, one of the purposes of this study was to identify specific gene for *M. abscessus subsp. bolletii* identification by the combination of several genes, and to build a powerful and reliable taxonomical tool for the MABC based on minimum and maximum identity values.

## MATERIALS AND METHODS

### Bacterial isolates

A total of 6 MABC isolates were obtained from different patients from the same hospital at the same time period (2009). Of them, 2 strains were isolated from sputum samples and 4 were obtained from Bronchoalveolar Lavage Fluid (BALF). All of the isolates had been classified as *M. abscessus* based on the results of biochemistry followed the Leao’s method[20].

### DNA Extraction

Bacterial DNA extraction was performed as described previously[21]. In brief, harvested the bacterium after growing in Middlebrook 7H9 liquid medium for 5 days, and then the samples were crushed with zirconia beads (1 mm in diameter) in a tissue disintegrator instrument. Total genomic DNA was extracted from the crushed suspension using a commercial ethanol precipitation kit according to the manufacturer’s instructions and stored at −20°C. Samples were sequenced on the Illumina HiSeq X10 sequencer, using 150 base-pair paired-end reads.

### Genome assemble

After checking the length and quality of the reads with FastQC (Version 0.11.8)[22]. The reads were de novo assembled with SPAdes(v3.13.0), using the ‘-careful’ setting and k-mers 21, 33, 55, 77, and 99[23, 24], and the assembled genome was manually trimmed the Short contigs(length than 3,000bp) and very low coverage contigs(less than 10).The result of the assembly was evaluated using QUAST (Version v.5.0.2, http://quast.bioinf.spbau.ru/;)[25].

### Vitro drug susceptibility testing

Using the microdilution method following the recommendations of the Clinical and Laboratory Standards Institute [Clinical and Laboratory Standards Institute (CLSI) of the rapidly growing mycobacteria with microdilution method][26]. The susceptibility results to ciprofloxacin, moxifloxacin and clarithromycin were judged by the established breakpoints from CLSI document (M24-A2-2011).

To evaluate the traditional identification taxonomy method, 16S rRNA (1,468-bp), *rpoB* (409-bp), *erm*(41)(GenBank accession number:CU458896.1), *erm*(42)(GenBank accession number: FJ358487.1) were selected for comparison[27, 28]. Nucleotide sequence were extracted from the reference sequences *M. abscessus subsp. abscessus* ATCC 19977 and *Mycobacterium_abscessus_subsp_bolletii* CIP 108541 using the traditional primers(The primer used in this study was listed in supplemental table 1). For comparision, the *erm*(41) fragment was selected with the same beginning and end with *erm*(42) fragment. The identities among the fragments were got by BLASTN searches with 1E-5 as a cut-off value [29]. And the fragments from the genomes were gotten by TBtools[30].

**Table 1.** ANI, GGDC, DDH, *rpoB* and AAI genomic relatedness among the type strains *M. abscessus subsp. abscessus* ATCC 19977, *M. abscessus subsp. massiliense* GO 06 and *M. abscessus subsp. bolletii* CIP 108541

| Genome 1 | Genome 2 | OrthoANI<br>value (%) | Original ANI<br>value (%) | GGDC<br>distance | Formula 2 |  | *rpoB*<br>similarity |
| --- | --- | --- | --- | --- | --- | --- | --- |
|  |  |  |  |  | Dista<br>nce | DDH<br>estimate |  |
| *Mycobacterium_abscessus_subsp_abscessus* | *Mycobacterium_abscessus_subsp_bolletii* | 97.3906 | 97.1484 | 0.026672229 | 0.0267 | 77.20% | 97.78 |
| *Mycobacterium_abscessus_subsp_abscessus* | *Mycobacterium_abscessus_subsp_massiliense* | 97.3751 | 97.2282 | 0.027495489 | 0.0275 | 76.60% | 98.067 |
| *Mycobacterium_abscessus_subsp_bolletii* | *Mycobacterium_abscessus_subsp_massiliense* | 96.94 | 96.7821 | 0.031488937 | 0.0315 | 73.40% |  |

| Genome 1 | Genome 2 | OrthoANI value (%) | Original ANI value (%) | GGDC distance | Formula 2 |  |
| --- | --- | --- | --- | --- | --- | --- |
|  |  |  |  |  | Distan | DDH |
|  |  |  |  |  | ce | estimate |
| Mycobacterium_abscessus_subsp_abscessus | Mycobacterium_abscessus_subsp_bolletii | 97.3906 | 97.1484 | 0.026672229 | **0.0267** | 77.20% |
| Mycobacterium_abscessus_subsp_abscessus | Mycobacterium_abscessus_subsp_massiliense | 97.3751 | 97.2282 | 0.027495489 | **0.0275** | 76.60% |
| Mycobacterium_abscessus_subsp_bolletii | Mycobacterium_abscessus_subsp_massiliense | 96.94 | 96.7821 | 0.031488937 | 0.0315 | 73.40% |

| SeqA | SeqB | AAI | SD | N | Omega | Fr<sub>x</sub> |
| --- | --- | --- | --- | --- | --- | --- |
| Mycobacterium_abscessus_subsp.massilien<br>se | Mycobacterium_abscessus_subsp.massilien<br>se | 100 | 0 | 455<br>6 | 4558 | 99.9561<br>2 |
| Mycobacterium_abscessus_subsp_abscessus | Mycobacterium_abscessus_subsp.massilien<br>se | 97.9505269<br>3 | 4.73846818<br>9 | 427<br>0 | 4558 | 93.6814<br>4 |
| Mycobacterium_abscessus_subsp_abscessus | Mycobacterium_abscessus_subsp_abscessus | 100 | 0 | 494<br>0 | 4942 | 99.9595<br>3 |
| Mycobacterium_abscessus_subsp_bolletii | Mycobacterium_abscessus_subsp.massilien<br>se | 97.6233239<br>1 | 4.93923620<br>4 | 424<br>5 | 4558 | 93.1329<br>5 |
| Mycobacterium_abscessus_subsp_bolletii | Mycobacterium_abscessus_subsp_abscessus | 97.6113439 | 6.81469740<br>2 | 435<br>3 | 4898 | 88.8730<br>1 |
| Mycobacterium_abscessus_subsp_bolletii | Mycobacterium_abscessus_subsp_bolletii | 100 | 0 | 489<br>2 | 4898 | 99.8775 |

The ANI values were gotten from http://enve-omics.ce.gatech.edu/g-matrix/.

In order to test method got in this study, all of the genomes (complete, chromosome, scaffold) were downloaded from NCBI genome (https://www.ncbi.nlm.nih.gov/genome/genomes/1360?). In order to evaluate the subspecies of MABC, average nucleotide identity (ANI), average amino acid identity (AAI) and genome to genome distance (GGDC) were calculated among the typical strains using KostasLab two-way average AAI calculator (http://enve-omics.ce.gatech.edu/aai/)[31], ANI (http://enve-omics.ce.gatech.edu/)[32], OrthoANI calculator[33], Genome-to-Genome Distance Calculator (http://ggdc.dsmz.de/ggdc.php)[34], fastANI[40] was also used to estimate the ANI among the genome datasets constructed in this study. In order to get the most effective gene to distinguish *M.bolleti* from other MABC, Up to date Bacterial Core Gene (UBCG) tool was chose to select gene from the 92 bacterial core genes[35]. The Phylogenetic tree was visualized using iTOL[36].

## Result

Patients were primarily male (4/6 patients) and greater than 65 years of age (4/6patients). Antimicrobial resistance patterns of *M. abscessus subsp. abscessus* and *M. abscessus subsp. massiliense* isolated from respiratory specimens were shown in supplemental table 2. The result confirmed the innate resistance of MABC to several drug classes, such as the cefoxitin. Fortunately, those isolates are still sensitive to clarithromycin in 3 days. And the isolate 1 was the only one with observed resistance to clarithromycin at 7 days. Genomes analyze result also showed that no known resistance-conferring mutation in the 23S rRNA was observed in those isolates.

Then study first verified that whether MABC belonged to the same specie or different species. The AAI, ANI, GGDC results of the typical isolates of the MABC were shown in table 1. Among the representative genome dataset as previously reported (unpublished result), using the full length *ropB* obtained from *M. abscessus subsp. abscessus* ATCC 19977 as reference, the similarities among MABC complex represent isolates were from 97.78%-100% (table 1 and figure 1). All the results showed MABC complex belonged to the same specie.

**Figure 1:**
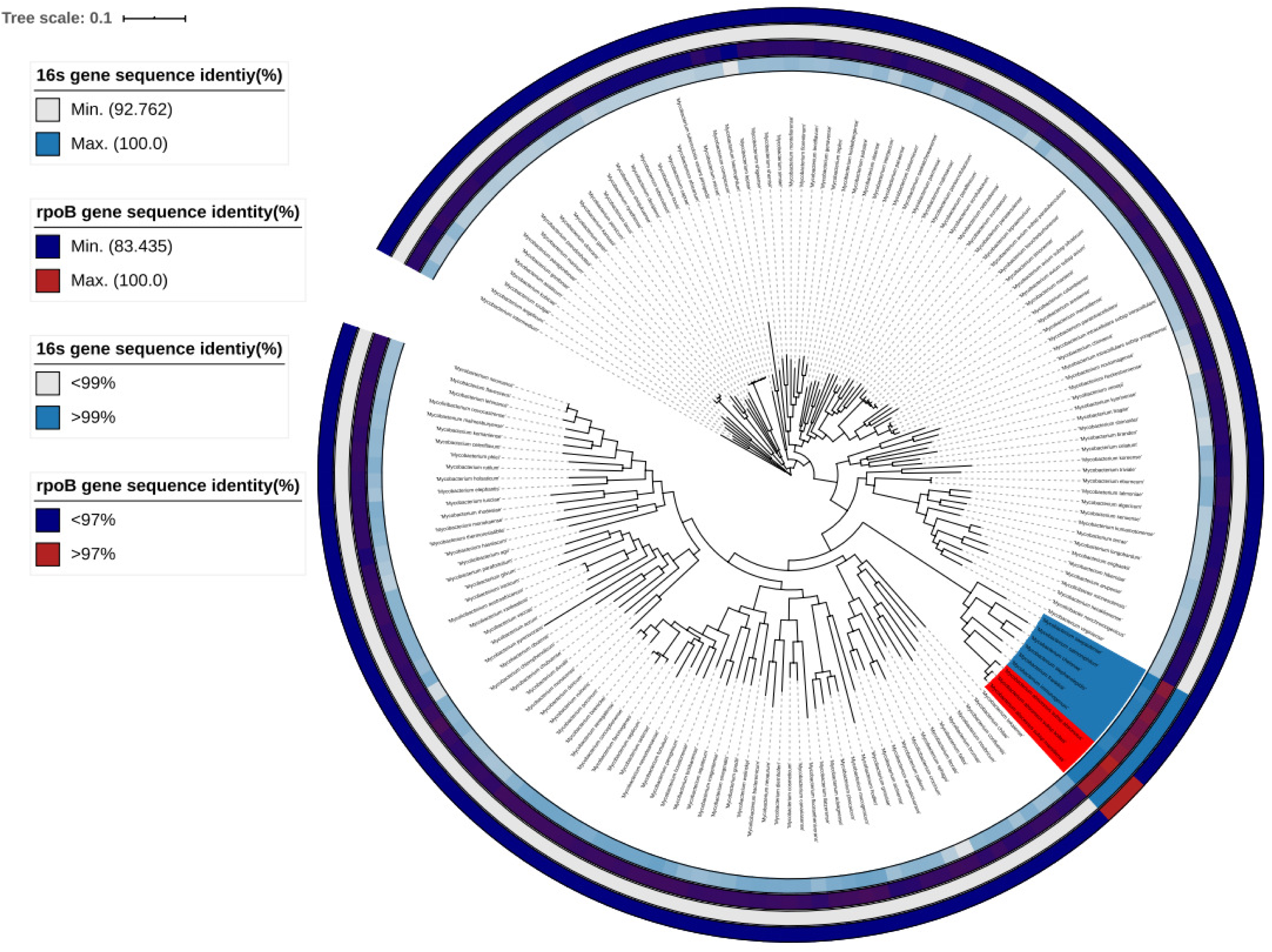
Phylogenetic relatedness of *Mycobacterium* spp and the identities of the 16s and *rpoB* genes. The inner range labeled with red and blue branches red branches correspond to the MABC, while the blue branches correspond to the neighbor species that were incorrectly identified as MABC by 16s identities. The grey and light blue range shows the identities of 16s rRNA and the the dark blue and red ranges show the identity of *rpoB*. The outside two ranges show the identities of 16s and rpoB genes with the cut-off values.

Then this study tested whether WGS comparison could further distinguish subspecies among the MABC complex. The ANI values for *Mycobacterium abscessus subsp massiliense* were from 96.72%-100% (Supplem table 6). And the ANI for *Mycobacterium abscessus subsp bolletii* were from 96.1263%-100% (Supplemental table 7). Because the overlap value of the ANI, the ANI value could not be used for subspecies distinguishing for MABC complex. The supplemental figure 1 also shows although the based on the ANI value could distinguish the species level but could not distinguish the sub species that are within the same species. Then another WGS based method-UBCG was used for subspecies verification. In accordance with our findings, based on the phylogenomic analyses of the UBCG (92 genes) of isolates from the public genome database showed the clear distance separating the three subsp of *M.abscessus* complex (figure 2 A). The genome which had been labeled the subsp names were with logo in figure. From the figure 2 A, we can see although many genomes were not labeled with supcies in the NCBI dataset, all the genomes with clear subspecies could be separate from other subspecies by different branches. So the UBCG is a good standard for subspecies verification. But using UBCG method for the subspecies classification took long time and with ambiguous boundaries (there is no clear cut-off value for UBCG method). So this study further tested the representative genes for species identification.

**Figure 2.**
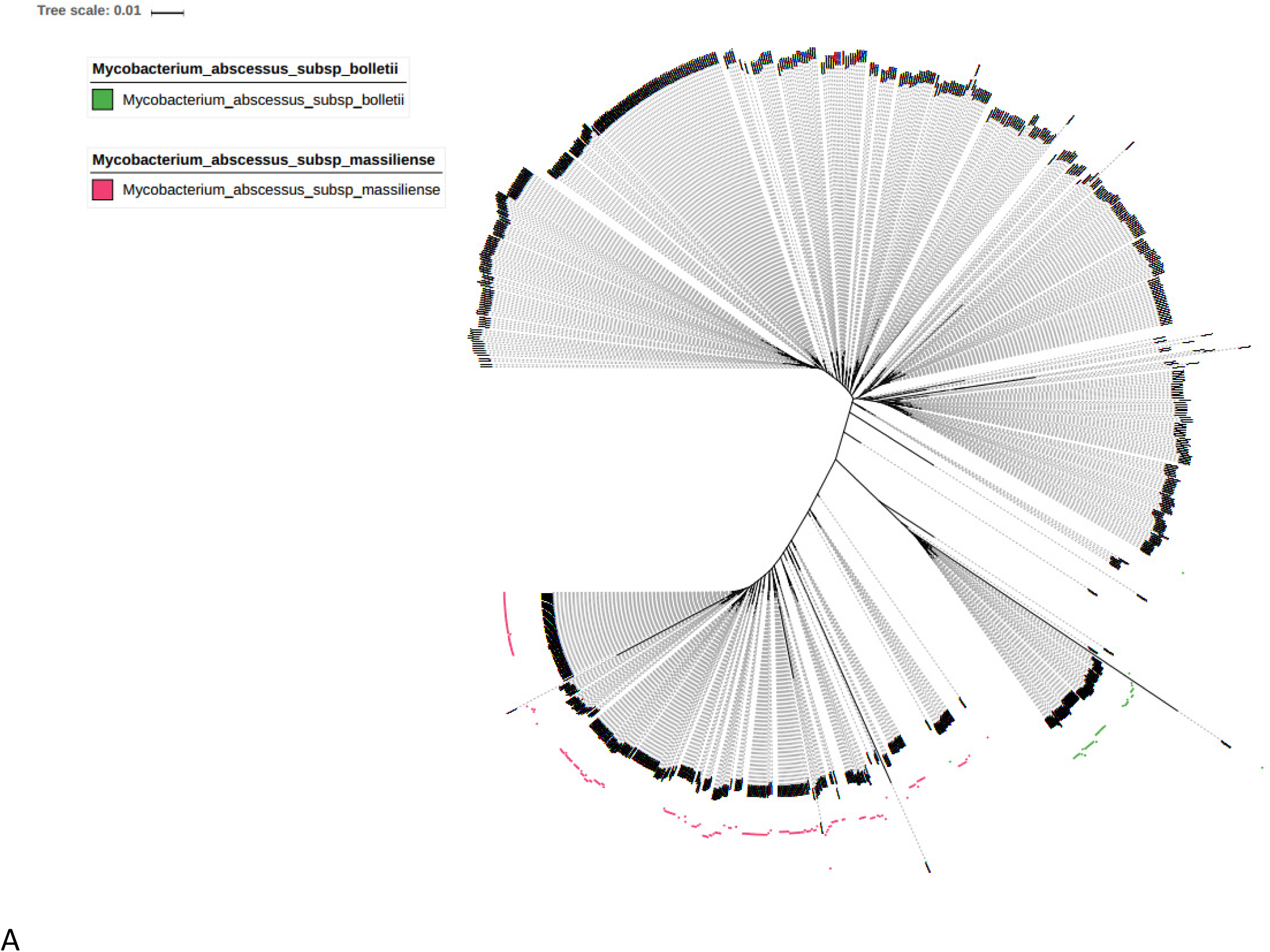
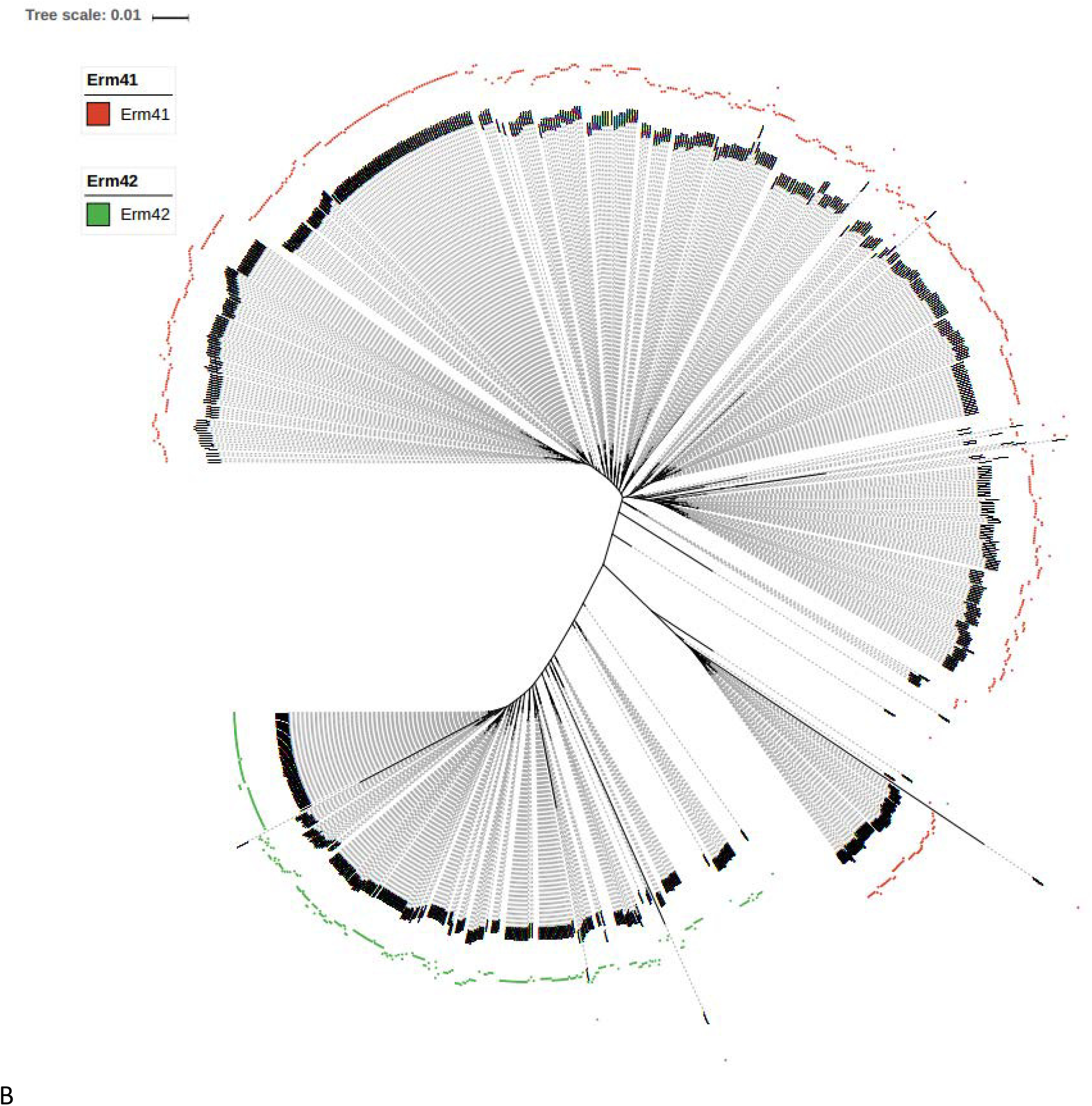
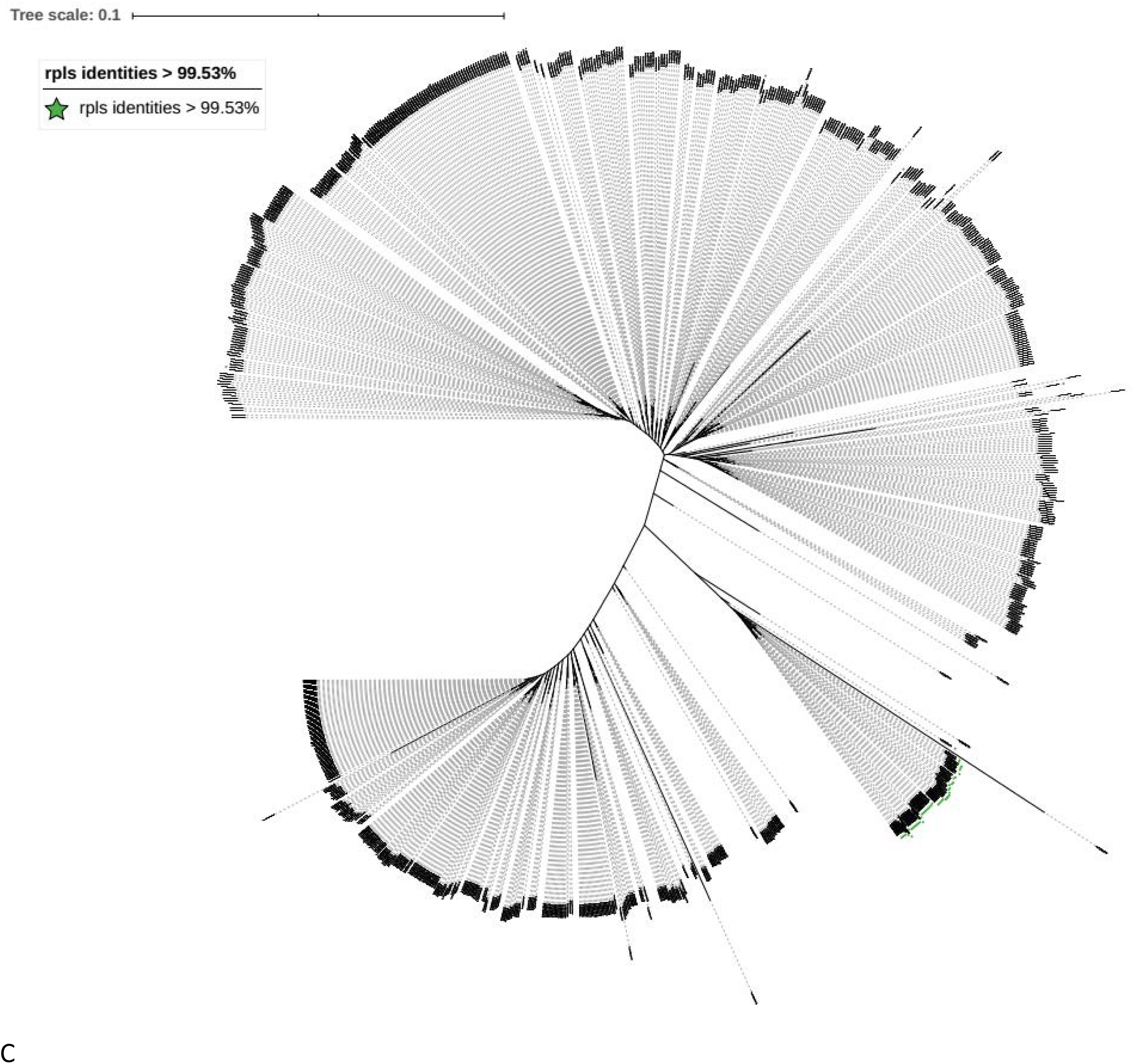
Phylogenetic tree for the genomes of the MABC complex. Phylogenetic tree inferred using UBCGs (concatenated alignment of 92 core genes) A Phylogenetic trees for MABC complex from the NCBI with all the clear label of *M. abscessus subsp. massiliense* and *M. abscessus subsp. Bolletii.* B Phylogenetic trees for MABC complex from the NCBI with all the clear label of the identities of *erm*(41) and *erm*(42). C Phylogenetic trees for MABC complex from the NCBI with *rpls* gene identities > 99.53% and *erm*(42) gene identities < 99.58%.

Firstly, in order to get the most accurate gene for identifying the MABC complex from other Mycobacterium spp, this study first evaluated the current used the gene fragment (Supplemental table 1). When we test the traditional marker for MABC identification, the *hsp* 65 and 16S–23S ITS primers could not be found in the genomes for the representative strains (Supplemental table 1). So this study only compared the 16s and *rpoB* genes. Then this study found that when using the fragment gotten by the primers rpoB-mycoF and rpoB-mycoR, the identity between the fragments from *M. abscessus subsp. abscessus* ATCC 19977 and *M. abscessus subsp. bolletii* CIP 108541 was 96.81. As the identity was below 97% which was settled by the traditional cut-off value, this study then used the full length rpoB genes for comparison. The similarities of the 16s and *rpoB* genes were shown in Figure 1. As shown in Figure 1, compared with 16s identity the rpoB genes identity could better distinguish the MABC with other Mycobcacterium spp. While 16s identities could not distinguish the MABC complex with similar species such as *Mycobacterium_salmoniphilum, Mycobacterium_immunogenum, Mycobacterium_stephanolepidis, Mycobacterium_franklinii, Mycobacterium_chelonae, Mycobacterium_saopaulense.* To verified whether the cut-off value could suit for all the MABC complex isolates, the genome dataset download from NCBI was used for testing. As the Supplemental_table 4 shows the *rpoB* identities were among 97.697%-100%. (893 isolates from 910 totals isolates, 7 of the genomes in the public databases were labeled contaminated were excluded from the dataset. Another 10 isolates in the genome database were shown in supplemental Supplemental_table 4, and all genome based identify method showed that they were very far from MABC complex). So the full length of *rpoB* gene and the cut-off value 97% could be used for MABC complex identification. Then this study tested whether *rpoB* could distinguish the subspecies. This study first utilized dataset of *M. abscessus subsp. Bolletii* from NCBI to test whether the full length *rpoB* gene could be used as subspecies marker. But when using the *rpoB* gene from *M. abscessus subsp. abscessus* ATCC 19977, the identities are ranging from 97.697-97.896%, and using the *rpoB* gene from *M. abscessus subsp. massiliense* the identities are from 98.052-99.742%. The *M. abscessus subsp. bolletii* CIP 108541 identities are ranged from 97.811 to 100%. As the ranges are overlapping with each other, using different rpoB genes could not distinguish other.

After using *rpoB* for MABC species identification, this study then verified the current used method for subspecies identificaiton. This study first selected the erythromycin ribosomal methylase gene, *erm*(41) for the further distinguishing the subspecies. The identities of *erm*(41) and *erm*(42) were shown in supplemental table 8. When using *erm*(41) fragment from *Mycobacterium abscessus subsp abscessu* ATCC 19977 as reference, the identities were from 97.893%-100% (586/893); and *erm*(42) fragment *Mycobacterium abscessus sub sp massiliense* (GenBank accession number: FJ358487.1) were 99.58%-100% (307/893). Because there were no genomes could share the identities with both *erm*(41) and *erm*(42), this study used identity 99.58% for *erm*(42) as the cut-off value for selecting *Mycobacterium abscessus sub sp massiliense* from the MABC complex. Then using already existed labeled *Mycobacterium abscessus subsp* tested the classification, and the results were shown in figure 2 B. From the figure 2 B we can see there were no cross between *erm*(41) and *erm*(42) (The genomes which contained both *erm*(41) and *erm*(42) are located in different branches). The figure showed that the *erm*(42) were exactly matched the *Mycobacterium abscessus subsp massiliense.* So does the *erm*(41) with *Mycobacterium abscessus subsp bolletii* and the isolates without special labeled. So the fragment of *erm*(42) with the cut-off value 99.58% was used for the identification of *Mycobacterium abscessus sub sp massiliense.*

In order to further distinguish the *Mycobacterium abscessus subsp abscessus* and *M. abscessus subsp. Bolletii*, this study first tested the traditional marker the fragment of *rpoB*. According to the traditional method *M. abscessus subsp. Bolletii* genomes should be group together according to identities of *rpoB* fragment. This study tested the identies of *rpoB* fragment from the *M. abscessus subsp. Bolletii* with the *rpoB* fragment from all the three species of MABC complex. The *rpoB* identities among the *M. abscessus subsp. Bolletii were* from 95.88% to 99.73 %, when using the fragment from *Mycobacterium abscessus subsp massiliense* GO 06. The identities were from 94.16% to 100%, when using the fragment from *Mycobacterium abscessus subsp abscessu* ATCC 19977, and the identies were from 95.88% to 100% when using the fragement from *M. abscessus subsp. bolletii* CIP 108541 (Supplemental table 9). As the *ropB* gene could not distinguish the *M. abscessus subsp. Bolletii*, this study tried to find a marker gene for *M. abscessus subsp. Bolletii* from the UBCG selected genes. All the 92 genes tested in UBCG were evaluated in this study. From all those individual gene trees, when using *rpls* gene for drawing the Phylogenetic tree, fewest genomes were involved with the genomes which had been labeled with *Mycobacterium abscessus subsp bolletii*(Figure 3 A). Then in order to get the cut-off value for the *rpls* gene for the *Mycobacterium abscessus subsp bolletii* identification, this study first used full length *rpls* gene identities(Supplemental table 10 A). Because two genomes’ identities were little far from other genomes, this study selected partial common *rpls* gene frgement share by *Mycobacterium abscessus subsp bolletii* for reference (Supplemental file 11). Although *Mycobacteroides abscessus subsp. bolletii* 50594 was labeled as *Mycobacteroides abscessus subsp. bolletii*, it carried *erm*(42) gene instead of *erm*(41). What is more this strain was also labeled as *Mycobacterium massiliense* 50594 (heterotypic synonym). So this study excluded this genome from *Mycobacteroides abscessus subsp. bolletii* dataset, and the cut-off value for *rpls* fragment identity was 99.53% (Supplemental table 10 B). Then study used the fragement and the cut-off value to test all the MABC complex genomes from the NCBI. The genomes with the identity of *rpls* fragement > 99.53% and identity of *erm*(42) fragement < 99.58% were exactly the branch of the *Mycobacteroides abscessus subsp. bolletii* (Figure 2 C, A and Supplemental table 10 C). So *rpls* gene fragment and the cut-off value 99.53% was selected for the *Mycobacterium abscessus subsp bolletii* identification.

**Figure 3.**
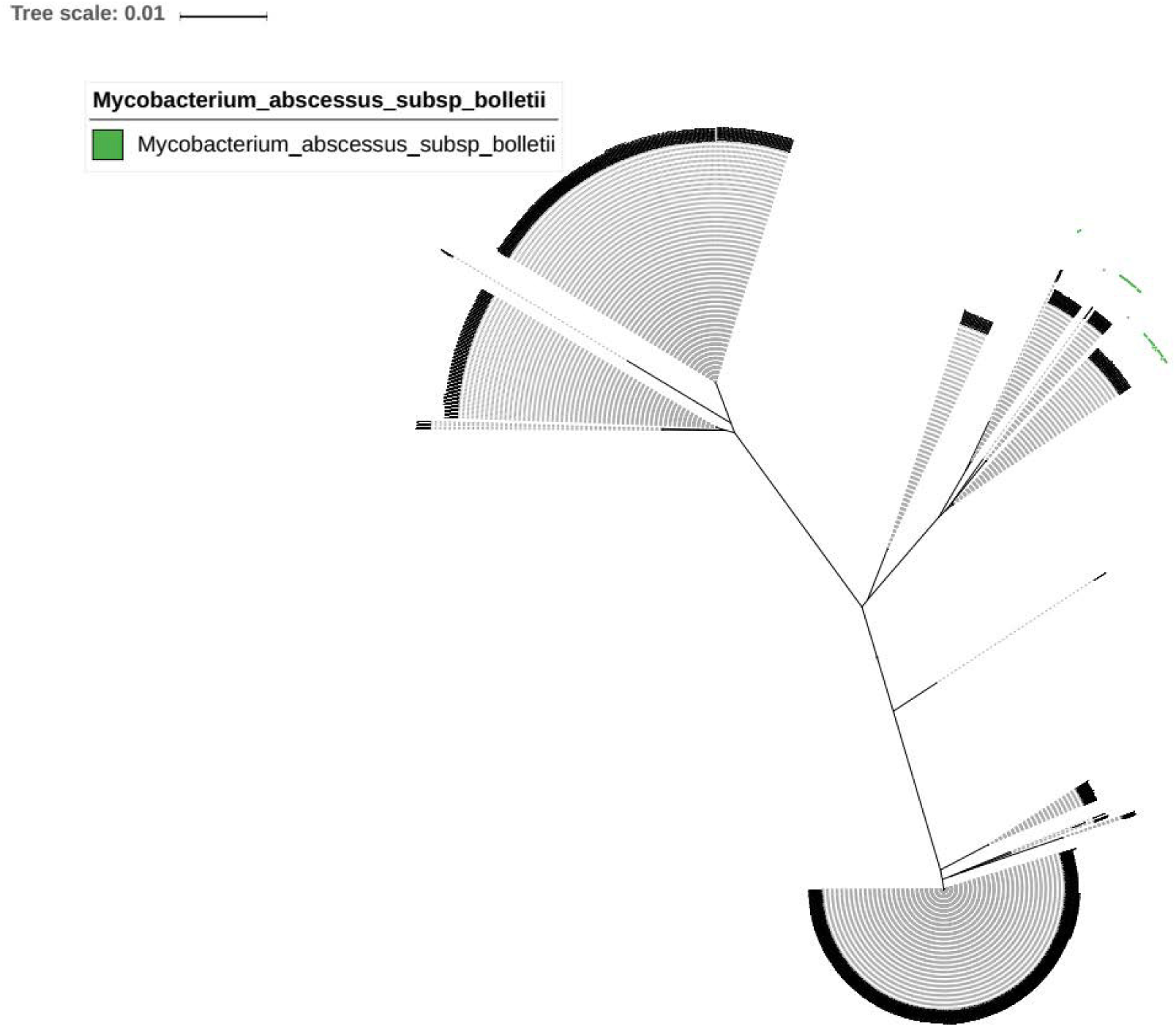
The phylogenomic tree for identification of *Mycobacterium abscessus subsp bolletii* A phylogenomic tree based on *rpls*. The green square labeled the *Mycobacterium abscessus subsp bolletii* genomes with clear labeling as *Mycobacterium abscessus subsp bolletii* in NCBI database.

The clinical isolates involved in this study were submitted to NCBI with the Accession number PRJNA594106. Last but not least, this study summary the analyzing data (Figure 4) and then contrsusted the software NucleotideQuery for the MABC complex classification (Attachment file 12) which is accurate and user-friendly (No need for installation, and could detect the genomes directly)..

**Figure 4.**
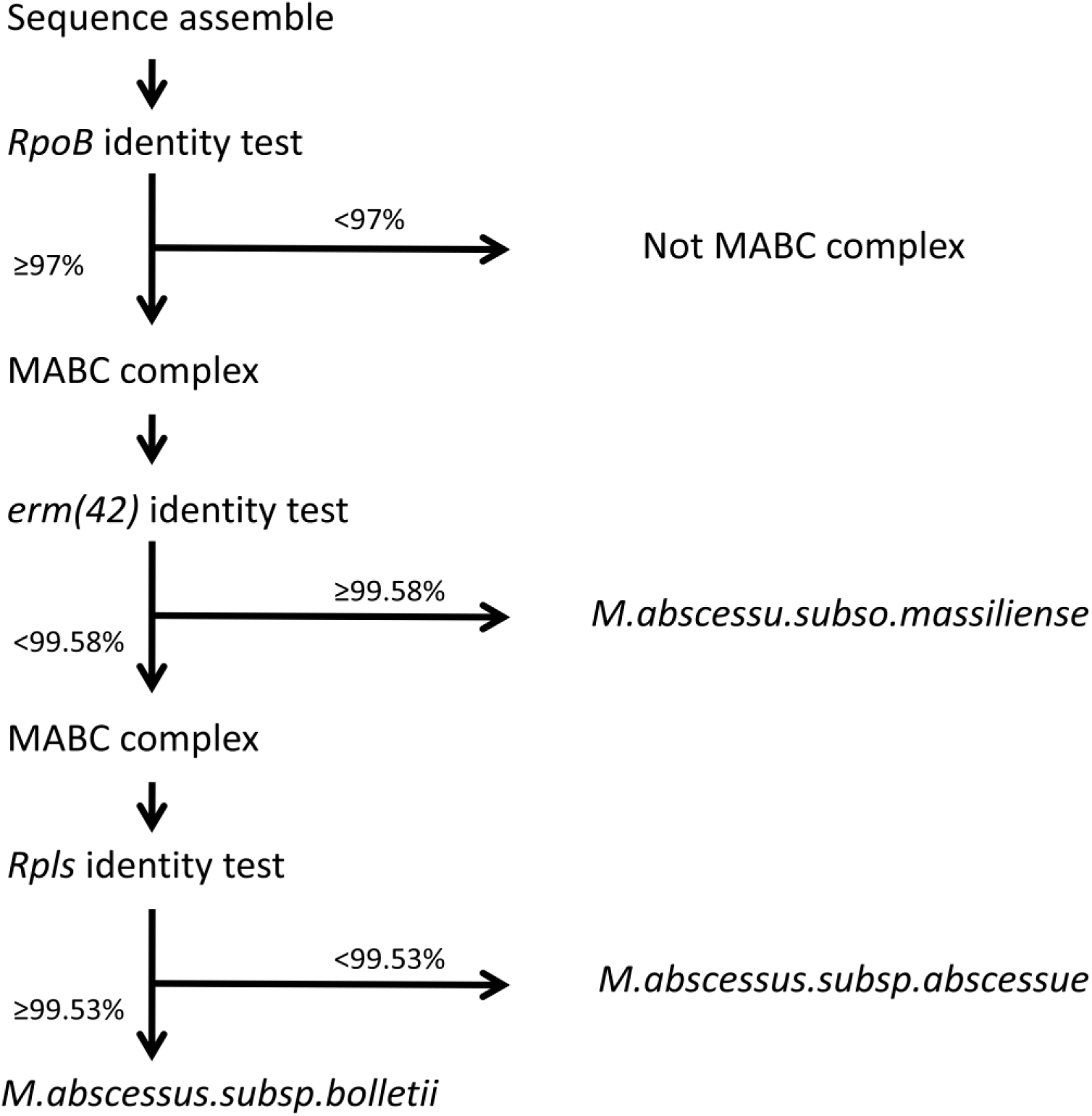
Flow chart of the study software.

## Discussion

All of the three *M.abscessus* subsp had been reported could cause serious infections such as pulmonary [37], and the controversies about MABC’s taxonomic status have never stopped. The proposed and generally accepted species boundary for AAI, ANI and dDDH values are 95%, 95∼96 and 70 %, respectively [34, 38, 39]. As the result showed in Table 1, the AAI values among the MABC typical strains were 97.61%-97.95%, and all larger than the cut-off value 95%. The ANI values among the MABC typical strains were 96.94%-97.39% (the results gotten by different softwares were similar as the Table 1 shown); and they were all larger than cut-off value 96%. the GGDC value among the MABC typical strains were 73.4%-77.2%. The above results of this study supported the current classification of *M. abscessus subsp. abscessus, M. abscessus subsp. massiliense* and *M. abscessus subsp as sub species* are belonged to the same specie.

The most common method to identify the MABC complex isolates were based on the *rpoB*, but this method had been questioned[40]. Andt he cut-off value of *rpoB* gene for MABC complex and even for the subsp in MABC complex had been studied for a long time[28, 41-43]. To our knowledge this study first time using the full length *rpoB* gene as the marker gene. What is more this study involved the largest number of MABC complex (all the available genomes in the NCBI genome). With the cut-off value of this study of *rpoB* to identifythe MABC complex was 97.697% that could distinguish the MABC complex with other *Mycoabcterium* spp. As showed in supplemental table 5, the three subspecies in MABC could not be distinguished with each other by *rpoB* only. This study then further evaluated the current used marker, and selected the 97.893 and 99.5588 for *erm*(41) and *erm*(42) fragement, respectively, which would distinguish *M.abscessus subsp. Massiliense* with *M.abscessus subsp.abscessus*. To our knowledge, this study is the first time to set the clearly cut-off value of the *erm* gene during the distinction of MABC complex. The Figure 2 showed that these cut-off values were suitable for *Mycobacterium abscessus subsp massiliense* identification.

Due to the clinical significance, all of the three subsp should be differentiated with each other rapidly. Differed with *M.abscesusssubsp.massiliense, M.abscessus subsp.bolletii* harbored a inducible functional erythromycin ribosome methyltransferase *erm*(41)[17]. Thus, this study further designed new target genes to distinguish *M.abscessus subsp.bolletii* from *M.abscessus subsp.abscessus*. Scientists had noticed the importance of identification of the subsp of MABC complex for a long time, but the previous reports for taxonomy of *Mycobacterium abscessus subsp massiliense* were ambiguous, especially for the cut-off value[44]. The same situation also occurs for the *Mycobacterium abscessus subsp bolletii*, previous reports used *rpoB, hsp65* or 16S–23S ITS for identification [44-48], and no cut-off value was set for each gene. Furthermore, the previous reported primers *hsp*65 and 16S–23S ITS[37]could not be found in the genomes for the two representative strains. So this study did not involved these genes for testing. In contast with the tradiotinal marker – the fragement of *rpoB* frangment[49, 50], this study also had showed that the *rpoB* gene could not distinguish *Mycobacterium abscessus subsp massiliense* with *Mycobacterium abscessus subsp bolletii*as shown in the attachment table 9. When using *rpoB* fragment from *M.abscessu.subsp.bolletii* as reference, the identities among the labeled *M.abscessu.subsp.bolletii* genomes were from 95.878% to 100%; when using *rpoB* fragment from *M.abscessu.subsp.massiliense* as reference, the identity among the labelled *M.abscessu.subsp.bolletii* genomes were from 95.878%-99.734%. As the ranges were over cross with each other, so this study believed the *rpoB* fragments could not be used for *M.abscessu.subsp.bolletii* identification. Previous study also confirmed that single gene was not good enough for subsp taxonomy[51]. Other methods like the pulsed-field gel electrophoresis are complex and costly that are not suitable for clinical field[52]. In order to find out the most effective gene for distinguishing *M.abscessus.subsp. bolletii*, all 92 genes selected by the UBCG were evaluated by the number of the smallest group including *M.abscessus.subsp. bolletii*. And then this study got the cut-off value by the *M.abscessus.subsp. bolletii.* After verified the selected gene *rpls* and cut-off value using the MABC complex dataset, this study demonstrated the *rpls* fragement with the identity cut-off value 99.53% could be served as the unique *M.abscessus.subsp. bolletii* identication marker.

The ANI has been validated as prokaryotic species taxonomy tool, where ANI values higher than 95–96% are consistent with strains belonging to the same species. To our knowledge, no cut-off has been proposed yet to define the boundary of subspecies. Although most of the *M.abscessus.subsp.abscessus* genomes were not labeled in the genome database, the ANI value could not distinguish *M.abscessus.subsp.bolletii* with *M.abscessus.subsp. massiliense*(attachment table 6 and 7). This study also showed that the ANI value was not suitable to distinguish the sub species of MABC complex. But when using WGS assemble and then analyzed by our software; this rapid approach offers the accurate, user-friendly MABC complex subspecies identification using *rpoB, erm(42)* and *rpls* are sufficiently reliable to serve as the routine methodology in clinical field. So, our method was with high clinical meaning. Conclusion

This study reported genome sequences and genomic features of 6 MABC isolates derived from one hospital in China. The results of this study showed that the distance among MABC complex was insufficient to warrant distinction at the species level, so there are subspecies in MABC complex. And after verified the taxonomy of MABC complex, we had developed an user-friendly accurate method based on a set of genes for differentiation at the subspecies level *M.abscessus subsp. massiliense, M.abscessu subsp.bolletti* and *M.abscessus subsp.abscess*. Using all available sequences to test the method, and the sequence analysis of public databases indicated that the combination by this design could be high discrimination among MABC complex.

Whole-genome sequencing technology has been more and more widely used in clinical diagnosis, developing more user-friendly software would greatly facilitate the acquisition of more-precise information about pathogen, to aid in the choice of more discriminate therapies.

## ACKNOWLEDGMENT

The authors thank the Dr Dong Yun for for critical reading of the manuscript and members of the Xinchun lab for useful discussion.

Tis work was supported by the National Natural Science Funds of China (81802060) and the start-up funding of Shenzhen University

